# Towards complete carbon utilization: Improved methane yield from formate and hydrogen co-feeding through constitutive formate dehydrogenase-gene expression in *Methanothermobacter thermautotrophicus* ΔH

**DOI:** 10.64898/2026.03.21.713158

**Authors:** Aaron Zipperle, Largus T. Angenent, Gerben R. Stouten, Bastian Molitor

**Author notes:** **Co-corresponding authors**, Gerben R. Stouten -; Bastian Molitor. **CRediT** Aaron Zipperle: Conceptualization, Data Curation, Formal Analysis, Investigation, Methodology, Visualization, Writing – original draft and review & editing Largus T. Angenent: Conceptualization, Funding Acquisition, Project Administration, Supervision, Writing – review & editing Gerben R. Stouten: Conceptualization, Data Curation, Formal Analysis, Investigation, Methodology, Visualization, Project Administration, Supervision, Validation, Writing – original draft and review & editing Bastian Molitor: Conceptualization, Funding Acquisition, Methodology, Project Administration, Supervision, Validation, Writing – review & editing.

## Abstract

Formate is emerging as an important molecule in carbon capture and utilization technologies. However, its low electron density makes formate less attractive for energy storage. Some hydrogenotrophic methanogens can reduce formate to methane, thereby upgrading it into an established energy carrier. The bottleneck in this process is that 75% of the carbon is lost as carbon dioxide, and achieving a complete formate-to-methane conversion requires co-feeding hydrogen. However, hydrogen-dependent genetic regulation of formate metabolism inhibits simultaneous formate and hydrogen utilization in hydrogenotrophic methanogens. Here, we compared the catalytic performance of the genetically modified strain *Methanothermobacter thermautotrophicus* ΔH (pFdh) with *M. thermautotrophicus* Z-245 by conducting continuous cultivation at different hydrogen concentrations. While *M. thermautotrophicus* Z-245 is a natural formatotroph, *M. thermautotrophicus* ΔH (pFdh) was engineered to enable formate utilization *via* episomal expression of a formate dehydrogenase-gene cassette.

We found that *M. thermautotrophicus* ΔH (pFdh) can simultaneously utilize formate and hydrogen. It continuously consumed formate at ≈ 0.1 mM dissolved hydrogen, enabling a 75.6% formate-to-methane conversion efficiency. *M. thermautotrophicus* Z-245 showed a declining formate consumption at ≈ 0.016 mM and only reached a maximum stable efficiency of 36.3%. These results suggest that *M. thermautotrophicus* ΔH (pFdh) is largely insensitive to hydrogen-induced genetic regulation; however, it still faces redox-related metabolic limitations at dissolved hydrogen concentrations above 0.4 mM. Overall, the findings reveal a potential strategy to circumvent hydrogen-induced regulation of formate metabolism and identify *M. thermautotrophicus* ΔH (pFdh) as a promising biocatalyst for formate-to-methane conversion.

## 1. Introduction

In the transition toward a circular bioeconomy, carbon dioxide (CO_2_) capture and conversion technologies provide a promising bridge between renewable electricity generation and carbon-based chemical industry. One option with these technologies is to generate formate, which is a safe and fully soluble one-carbon (C1) compound (Hadi et al., 2025; Yishai et al., 2016). However, current processes that generate formate from renewable electricity and CO_2_, typically produce an aqueous solution at concentrations below 100 mM (Fernández-Caso et al., 2023; Quintana-Gómez et al., 2021; Sahoo et al., 2021). This means further up-concentration is needed for transportation, making it an ineffective energy carrier. Direct co-localized conversion is possible with biological processes, which are fully compatible with aqueous formate solutions and can use it as a substrate to produce compounds like mevalonate and isoprenol (Cowan et al., 2025). Nonetheless, formate remains a sub-optimal substrate because of its low energy density (Hanselmann, 1991). Therefore, further reducing formate into methane (CH_4_) might be a better solution to close the carbon cycle and create an energy-dense and infrastructure-compatible energy carrier. Biotechnologically reducing CO_2_ to CH_4_ is an established process that uses molecular hydrogen (H_2_) as an electron donor (Angenent et al., 2022). The same principle could be applied to reduce formate into CH_4_, thereby, storing renewable energy.

Many methanogens can use formate as a substrate in defined mineral media (Tejedor-Sanz et al., 2024). These properties make methanogens suitable candidates to convert formate to CH_4_. Nonetheless, formate disproportionation only yields 25 mol-% CH_4_ (on a carbon basis), the remainder is oxidized back to CO_2_. Co-feeding molecular H_2_ would fully reduce formate-derived CO_2_ to CH_4_ in a single bioprocess. However, elevated H_2_ levels suppress formate utilization in methanogens, which is a phenomenon that studies have linked to transcriptional control of formate dehydrogenase (*fdh*)-genes (Costa et al., 2013; Hendrickson et al., 2007; Nölling & Reeve, 1997). Thus, currently a full carbon conversion of formate to CH_4_ is not possible.

Here, we investigated whether decoupling *fdh*-gene expression from endogenous regulation would improve process stability and carbon utilization in formate-driven methanogenesis under H_2_ exposure. To test this, we co-fed formate and H_2_ to *Methanothermobacter thermautotrophicus* ΔH pMVS1111A:PhmtB-fdh_Z-245_, hereafter referred to as *M. thermautotrophicus* ΔH (pFdh) (Fink et al., 2021). For this strain, an *fdh*-encoding gene cassette from the strain *Methanothermobacter thermautotrophicus* Z-245 (*i*.*e*., a strain that natively grows with formate as the sole electron and carbon source) is expressed constitutively from a plasmid. This modification enables *M. thermautotrophicus* ΔH (pFdh) to use formate as a sole electron and carbon source, while the wild-type strain, *M. thermautotrophicus* ΔH, cannot natively grow with formate (Casini et al., 2023). We hypothesized that *M. thermautotrophicus* ΔH (pFdh) would maintain formate oxidation activity and support co-utilization of formate and H_2_ even at elevated H_2_ partial pressures by bypassing native transcriptional control of the *fdh*-gene expression. To test this hypothesis, we compared *M. thermautotrophicus* ΔH (pFdh) with the genetically inaccessible donor strain *M. thermautotrophicus* Z-245 across a gradient of H_2_ availability.

We cultivated both strains in chemostat bioreactors supplied with formate and subjected them to stepwise increases in H_2_ partial pressure. Chemostat cultivation is essential for disentangling the effects of gaseous substrates on methanogen metabolism, because it allows independent control of dilution rate, gas composition, and substrate supply, while maintaining the cultures at metabolic steady state. In this setup, we can quantify dissolved gas concentrations in the liquid phase and keep them approximately constant over time, which enables quantitative assessment of carbon balances, yields, and biomass specific rates. Therefore, bioreactor cultivation enabled us to directly assess whether constitutive *fdh*-gene expression improves carbon utilization and CH_4_ production under H_2_ exposure.

## 2. Materials and Methods

### 2.1 Strains and Media

We obtained *Methanothermobacter thermautotrophicus* Z-245 (DSM 3720) and *Methanothermobacter thermautotrophicus* ΔH (DSM 1053) from DSMZ (Braunschweig, Germany), and the strain *M. thermautotrophicus* ΔH (pFdh), which is harboring the plasmid pMVS1111A:PhmtB-fdh_Z-245_, from Fink et al., 2021. Both precultures and bioreactor cultures were grown in modified mineral salt medium (Balch et al., 1979) (**Supplementary Table S1**). Per liter, the medium contained: NaHCO_2_, 13.6 g; NH_4_Cl, 0.5 g; L-cysteine HCL, 0.5 g; NaCl, 0.45 g; KH_2_PO4, 0.23 g; K_2_HPO_4_, 0.17 g; MgCl_2_·6 H_2_O, 0.08 g; CaCl_2_·2 H_2_O, 0.06 g; Na_2_MoO_4_·2 H_2_O, 2.42 mg; Na_2_SeO_3_, 0.173 mg; Resazurin, 4 mL (0.025 weight-%); (NH_4_)2Ni(SO_4_)_2_, 1 mL (0.2 weight-%); FeCl_2_·4 H_2_O, 1 mL (0.2 weight-% ); trace element solution, 1 mL. Additionally, 0.06 mL Antifoam 204 (Sigma Aldrich) was added in bioreactor medium. For experiments in serum bottles, no antifoam was added and 6.0 g L^-1^ NaHCO_3_ was used as pH buffer. The addition of the 0.5 g L^-1^ L-cysteine-HCl, for reduction of the system, was different between bioreactors and pre-cultures, and is described below. The composition of the trace element solution is given in **Supplementary Table S2** (Casini et al., 2023).

### 2.2 Pre-cultures

Precultures for inoculation of the bioreactors were grown in 100-mL serum bottles containing 20 mL medium at 60°C in a shaking incubator (ISS-7100R, JEIO TECH, Republic of Korea). The medium was made anaerobic by flowing with N_2_/CO_2_ (80:20 vol-%) for 1 h and transferred into an anaerobic chamber (UNIlab Pro, M. BRAUN INERTGAS-SYSTEME GMBH, Germany). Inside the chamber, L-cysteine-HCl was added to the final concentration specified above, and the pH was adjusted to 7.2 with HCl. Serum bottles were sealed with butyl rubber stoppers and aluminum crimp caps and removed from the chamber. We then exchanged the headspace with 0.5 bar overpressure N_2_/CO_2_ (80:20 vol-%) and autoclaved the bottles, which were then stored at room temperature until use.

### 2.3. Bioreactor operation

Two 1.2 L jacketed glass bioreactors (Duran, Germany) were operated aseptically in parallel as chemostats for 20 days, with a working volume of 0.8 L and a hydraulic retention time (HRT) of 2 days, corresponding to a dilution rate of 0.5 day^-1^ (Klask et al., 2020). The temperature was maintained at 60°C using a recirculation thermostat water bath (KISS 104A, Peter Huber Kältemaschinenbau SE, Germany), and cultures were mixed with a cross-shaped stirring bar that was rotated at 160 rpm by a magnetic stir-plate (MIXdrive 6HT, 2mag AG, Germany). The pH was controlled at 7.2 with 3 M NaOH and 3 M H_3_PO_4_. Medium inflow and effluent outflow were supplied by peristaltic pumps (Watson Marlow, United Kingdom), and the inlet gas was delivered at 25 NmL min^-1^ using mass flow controllers (Alicat MC Series, USA). Inlet gas and off-gas compositions were monitored continuously by online mass spectrometry (Prima BT, Thermo Fischer Scientific, UK).

Autoclaved bioreactors containing medium were first sparged with N_2_ for 2 h, after which 50x concentrated autoclaved L-cysteine-HCl solution was added to further remove molecular oxygen (O_2_) from the broth. The same procedure was performed for media bottles. Bioreactors were inoculated with 20 mL of preculture and operated in batch without gas sparging until cultures reached an optical density at 600 nm (OD_600_) of approximately 0.1, at which point continuous chemostat operation was initiated. The H_2_ partial pressure of the inlet gas was increased stepwise in the following order: 0%, 0.3%, 1%, 3%, 10%, 30%, 90% (vol-%, balance N_2_) Each condition was maintained for at least one day before changing to the next gas composition.

After completing the stepwise H_2_ increases, both bioreactors were modified to enhance the gas-liquid mass transfer by implementing gas recirculation. The bioreactor headspace was connected to the inlet gas line *via* a diaphragm pump (NMP830, KNF, Germany), which recirculated the headspace gas through the microsparger. This increased the gross gas flow rate through the bioreactor to approximately 1.5 NL min^1^, while the net gas supply to the system remained at 25 NmL min^-1^. With these modifications, the bioreactors were operated at 30 vol-% H_2_.

### 2.4 Sampling and analysis

Bioreactor cultures were sampled at least once per day. Before each sampling, a 5 mL aliquot was withdrawn and discarded to flush the sampling line; a second 5 mL sample was then collected for offline analysis. The OD_600_ was measured using a spectrophotometer (BioMateTM160, Thermo Scientific, United States), and the pH was verified with a benchtop pH meter (pH1100L, VWR; LE422, Mettler Toledo, United States). Formate concentrations in the culture supernatant were quantified by HPLC (HPLC: SIL-40C with a RID-20A detector, Shimadzu Europa, Germany; Column: Aminex HPX-87H, 300 by 7.8 mm; Bio-Rad, USA), as described before (Casini et al., 2023). Furthermore, the bioreactors were checked weekly for cross-contamination by PCR amplification of strain-specific genes (**Supplementary Table S3**). In detail, 300 µL sample was heated for 13 min at 100°C (ThermoMixer® 460-0223, Eppendorf, Germany) and then cooled on ice. From this, 1 µL was amplified in a thermocycler (Mastercycler® pro S, Eppendorf, Germany) with Phire Plant Polymerase (Phire Plant Direct PCR Master Mix, Thermo Fischer Scientific, Dreieich, Germany) according to the manufacturer’s instructions.

### 2.5. Determining the volumetric gas evolution rates

Volumetric gas evolution rates r_i_ (mmol L^-1^ h^-1^) were calculated from the inlet gas flow rate, measured gas composition, and an inert molecular nitrogen (N_2_) balance. The molar gas outflow rate Q_out_ was determined from the known molar inlet gas flow rate Q_in_, and measured mole fractions of N_2_ in the inlet and outlet gas streams (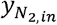 *and* 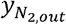 respectively):

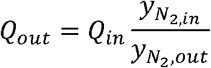

For each gas component *i*, the net evolution rate ER_i_ (mmol h^-1^) was then obtained from the gas balance:

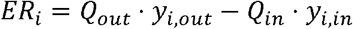

where *y*_*i,in*_ and *y*_*i,out*_ are the mole fractions of the compound *i* in the inlet and outlet gas, respectively. The volumetric gas evolution rate was calculated by normalizing to the bioreactor working volume V_L_:

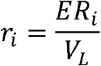

Biomass-specific uptake and production rates q_i_ (mmol_i_ gDW^-1^ h^-1^) were obtained from grams cell dry weight (gDW) using the measured OD_600_ and an *M. thermautotrophicus* specific conversion factor of 0.425 g L^-1^ per unit of OD_600_ (de Poorter et al., 2007).

### 2.6. Estimation of dissolved H_2_ concentration in the bioreactor

The availability of H_2_ to the microbes is governed by the dissolved H_2_ concentration in the liquid phase (cH_2_), which results from a balance between gas-liquid mass transfer and microbial uptake. At steady state in a chemostat, the H_2_ transfer rate (HTR) equals the H_2_ uptake rate (HUR), such that:

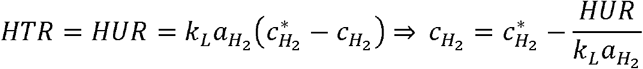

where 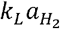 is the volumetric mass transfer coefficient for H_2_, 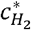 is the equilibrium dissolved H_2_ concentration at the applied gas partial pressure, and HUR is the biological H_2_ uptake rate determined from off-gas analysis (section 2.5 “Determining volumetric gas evolution rates”).

The volumetric mass transfer coefficient for H_2_ was estimated from O_2_ transfer experiments conducted under operating conditions (0.8 L broth, 25 mL min^-1^ gas flow, microsparger, 160 rpm agitation, 60°C).

Using a dynamic gassing-in/out method with N_2_ and air, we determined 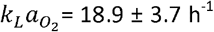. Reported diffusivities in water at 25°C are 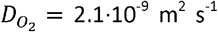 and 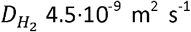 (Ferrell & Himmelblau, 1967), and their ratio remains approximately constant at 60°C (Cussler 2011). Assuming a surface-renewal dependence of *k*_*L*_*a* ∝ *D*^*n*^ with *n* between 0.5 and 0.67 (Danckwerts, 1951), this yields 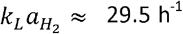. For operation with gas recirculation, 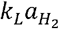 was extrapolated based on measurements at higher gas flow rates, giving an approximate value of 290 h^-1^. Further details of these calculations are provided in the Supplementary Materials. The equilibrium dissolved H_2_ concentration was obtained from Henry’s law, 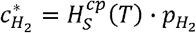, using a solubility-based Henry constant for H_2_ of 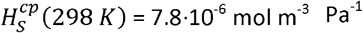 and a temperature correction factor C = 530 K (Sander, 2023). At 60°C this corresponds to 6.5·10^-6^ mol m^-3^ Pa^-1^, giving a maximum solubility of 0.65 mM (1.3 mg L^-1^ bar^-1^).

## 3. Results

To assess the effect of H_2_ co-feeding, *M. thermautotrophicus* Z-245 and *M. thermautotrophicus* ΔH (pFdh) were cultivated in chemostats supplied with formate and exposed to a stepwise increase in H_2_ partial pressures, ranging from 0 to 90 vol-% in the inlet gas (**Figure 1**). For each cultivation period, we quantified gas evolution and uptake rates, and estimated the maximum H_2_ transfer rate (HTR_max_) and the corresponding steady-state cH_2_ (**Supplementary Figure S1**). These calculations enabled us to distinguish between H_2_-limited and H_2_-excess regimes, and to assess how H_2_ co-feeding affected formate consumption and overall carbon utilization.

**Figure 1.**
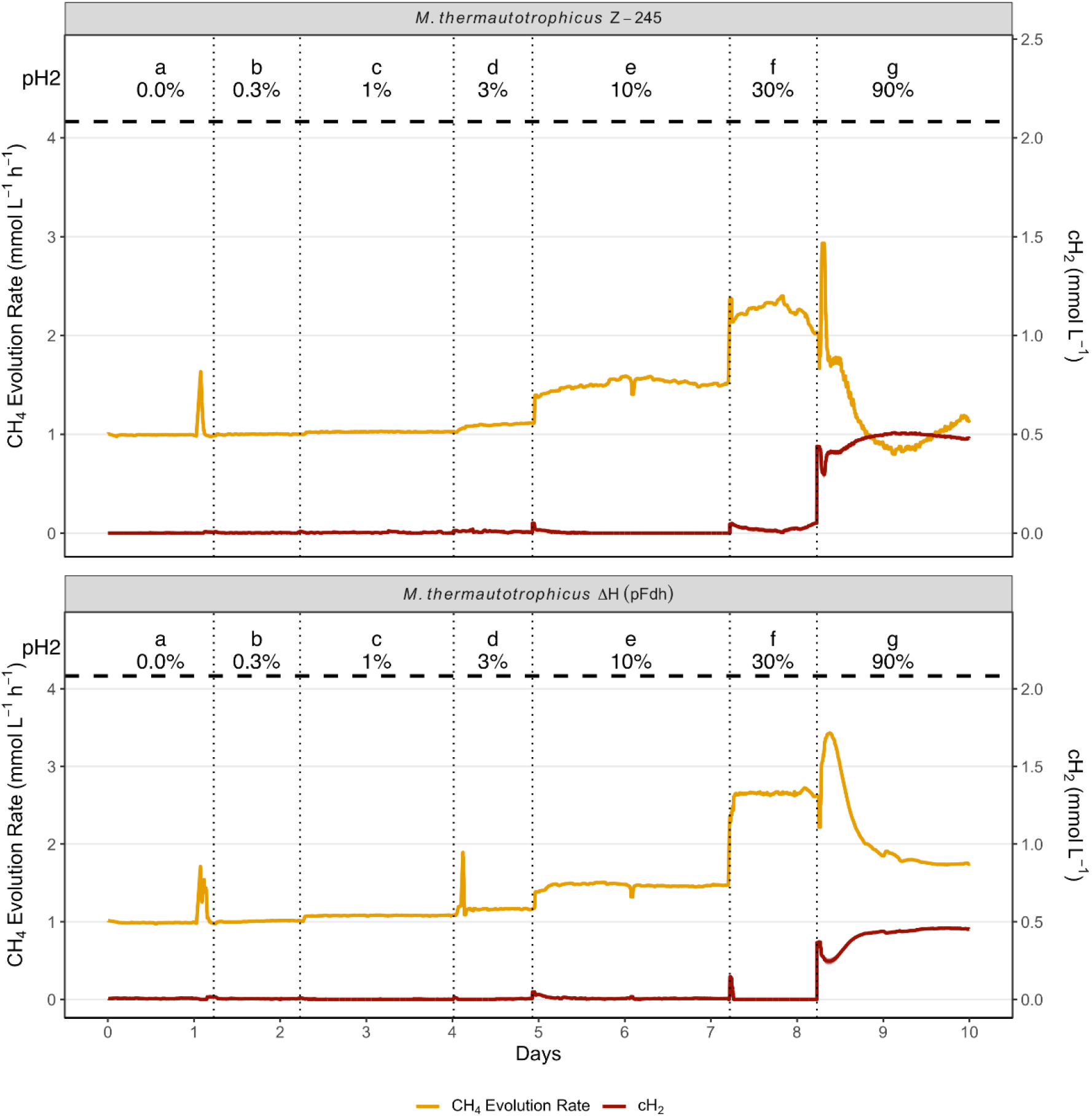
Methane evolution and dissolved H_2_ concentrations at increasing H_2_ partial pressure (pH_2_). M. thermautotrophicus Z-245 and M. thermautotrophicus ΔH (pFdh) were cultivated in a chemostat bioreactor with 4.17 mmol L ^-1^h ^-1^formate feed (black dashed line) and an increase in H_2_ partial pressure (pH_2_) in 7 steps (period a to g). From period b to e both strains consumed H_2_ at similar rate to which it was fed, keeping the dissolved H_2_ concentration in the liquid phase (cH_2_) close to 0 mmol L^-1^. A switc to 30 vol-% pH_2_ (period f) affected the formate consumption and methanogenesis in M. thermautotrophicus Z-245. Further increasing the pH_2_ to 90 vol-% caused a drastic increase in cH_2_ and both strains showed decreased formate uptake and methane evolution rates. Gray vertical bars indicate the transition point between the periods.

### 3.1. Influence of H_2_ partial pressures on formate consumption and carbon utilization

At low H_2_ partial pressures, where the majority of the reducing equivalents were supplied as formate, both strains behaved similarly. In these conditions, CH_4_ and CO_2_ evolution rates were comparable between the strains (**Figure 1 – periods a to e**). The H_2_ uptake rate matched the HTR_max_, indicating that the cH_2_ remained close to 0 mM and that H_2_ was a limiting substrate. Under these H_2_-limited conditions, both strains co-utilized formate and H_2_ without an observable negative effect on formate metabolism. However, when the strains were cultivated on formate alone in period a, *M. thermautotrophicus* ΔH (pFdh) produced quantifiable amounts of H_2_, which was not observed for *M. thermautotrophicus* Z-245 (1100 ppm *vs*. 200 ppm; **Supplementary Table S4** – **period a**). This effect was not observed in subsequent periods.

The first difference in CH_4_ and CO_2_ evolution between the strains emerged when the HTR_max_ reached similar amounts of reducing equivalents per hour as those from the formate supply (**Figure 1, Table 1** – **period f**). The H_2_ uptake rate of *M. thermautotrophicus* Z-245 did not match the HTR_max_, resulting in an increase in the dissolved H_2_ concentration to approximately 0.02 mM. The increase in cH_2_ was accompanied by reduced formate uptake and CH_4_ evolution rates, with downward trends throughout this cultivation period (**Figure 2**). These observations indicate that the bioreactor was no longer at a true steady state and that the balance between formate oxidation and methanogenesis was disturbed in *M. thermautotrophicus* Z-245 at 30 vol-% H_2_.

**Table 1.** Summary of main fermentation parameters for 4 cultivation periods with the highest H_2_ partial pressure (pH_2_). Data for M. thermautotrophicus Z-245 are shown in white and for M. thermautotrophicus ΔH (pFdh) in grey. Cultivation periods (for details see **Figure 1**) differ in the supplied pH_2_. Period h has a 10-fold higher gas mass transfer rate compared to f, while keeping the same pH_2_. Values represent period averages (∼24 h) except for period g, where the last 2 h of the period were used. Standard deviations are within 10%, except for HTR_max_, OD, Y_X/CH4_, and cultivation period h, which have a standard deviation within 27%. Data of all cultivation periods with standard deviations are provided in Supplementary Table S4. HTR_max_: maximum H_2_ transfer rate; HUR: H_2_ uptake rate; CER: CO_2_ evolution rate; MER: CH_4_ evolution rate; FUR: formate uptake rate; cH_2_: dissolved H_2_ concentration in the liquid phase; CU: carbon utilization; ē balance: the percentage of electrons recovered in measured products. In period g, no biomass measurement was taken in the time used for the calculation of the average, and therefore, the data is marked as not available (NA).

| Cultivation period |  | e |  | f |  | g |  | h |  |
| --- | --- | --- | --- | --- | --- | --- | --- | --- | --- |
| $pH_2$ | | 10 vol-% | | 30 vol-% | | 90 vol-% | | 30 vol-% (+ gas recycle) | |
| $HTR_{max}$ | mmol L<sup>-1</sup> h<sup>-1</sup> | 1.91 | | 5.74 | | 17.22 | | 56.66 | |
| HUR | mmol L<sup>-1</sup> h<sup>-1</sup> | 2.08 | 1.76 | 5.23 | 6.57 | 2.91 | 3.75 | - | 8.73 |
| CER | mmol L<sup>-1</sup> h<sup>-1</sup> | 2.48 | 2.62 | 1.64 | 1.45 | 0.42 | 1.50 | - | 0.76 |
| MER | mmol L<sup>-1</sup> h<sup>-1</sup> | 1.52 | 1.46 | 2.3 | 2.64 | 1.04 | 1.74 | - | 3.16 |
| FUR* | mmol L<sup>-1</sup> h<sup>-1</sup> | 4.00 | 4.07 | 3.49 | 4.04 | 1.58 | 3.24 | - | 3.91 |
| OD | - | 0.22 | 0.21 | 0.3 | 0.38 | 0.23 | 0.34 | - | 0.34 |
| $cH_2$ | mM | 0.00 | 0.01 | 0.02 | 0.00 | 0.48 | 0.46 | - | 0.10 |
| $Y_{X/CH_4}$ | gDW mol<sup>-1</sup> | 1.30 | 1.26 | 1.14 | 1.28 | NA | NA | - | 0.95 |
| CU | % | 36.4 | 35.0 | 65.9 | 63.4 | 73.3 | 57.1 | - | 75.8 |
| $e^-$ balance** | % | 101.3 | 102.3 | 98.4 | 95.9 | 96.6 | 96.0 | - | 94.7 |

\* Formate uptake rate as the sum of $CH_4$ and $CO_2$ evolution, discounting biomass formation.
\*\* Disregarding biomass

**Figure 2.**
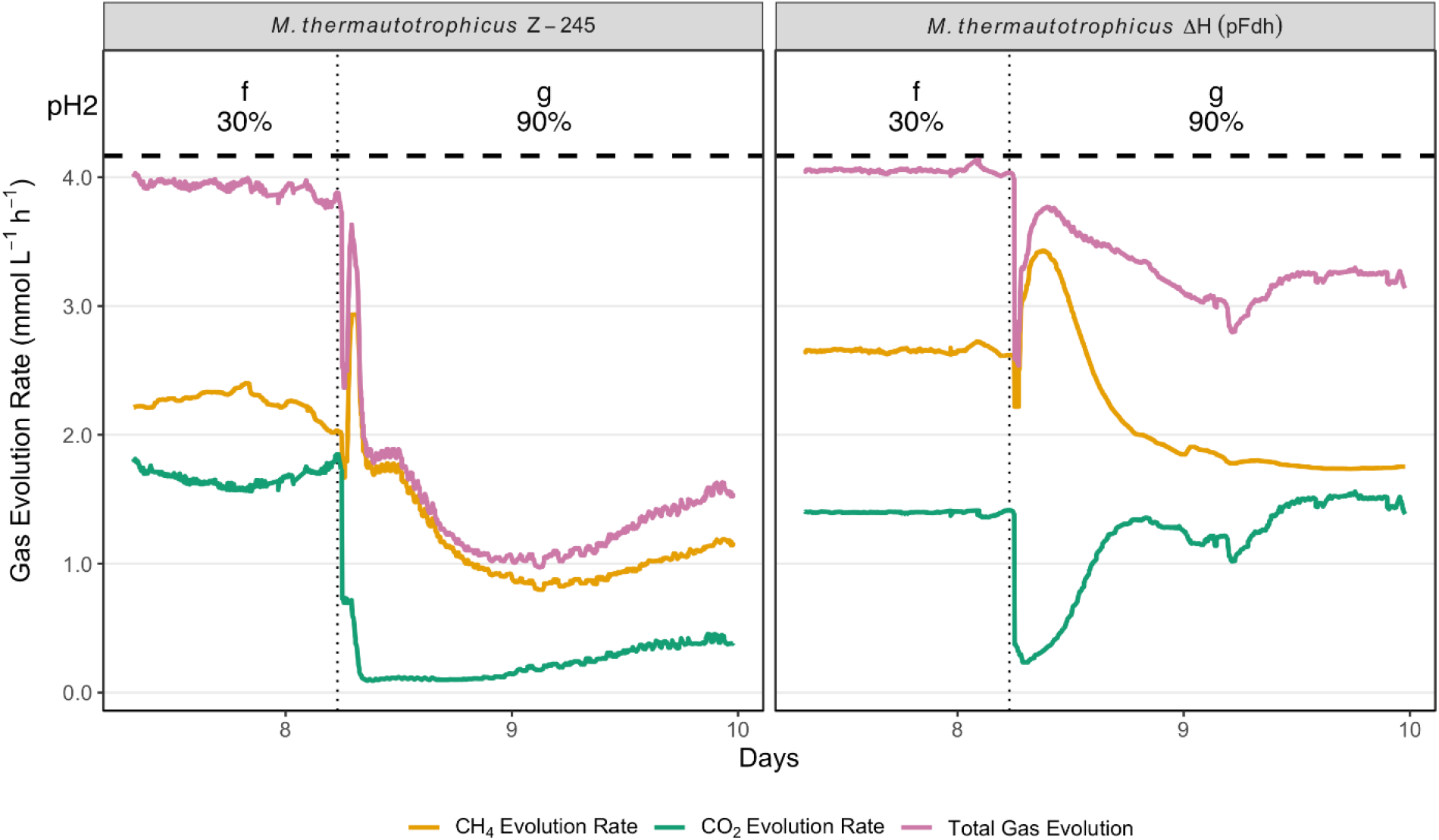
Gas evolution rates for M. thermautotrophicus Z-245 and M. thermautotrophicus ΔH (pFdh) during formate and H_2_ co-feeding for the switch from cultivation period f to g (black dotted line). We increased the H_2_ fraction in the inflow from 30 vol-% to 90 vol-%. Formate was fed at 4.16 mmol L^−1^ h^−1^ reflecting the carbon in flux (horizontal black dashed line). The sum of CO_2_ an CH_4_ evolution are shown in violet, and represent the formate uptake rate, disregarding biomass production. The measure formate concentration in the liquid phase is shown in Supplementary Figure S3.

In contrast, *M. thermautotrophicus* ΔH (pFdh) remained stable under the same conditions. Formate uptake was unaffected by the increased H_2_ supply, and CH_4_ evolution was stable at 2.64 ± 0.02 mmol L^-1^ h^-1^, corresponding to a carbon utilization of 63.4 ± 0.4 mol-% (**Table 1**). Thus, constitutive expression of the *fdh* cassette enabled *M. thermautotrophicus* ΔH (pFdh) to maintain stable co-utilization of formate and H_2_ in a regime in which *M. thermautotrophicus* Z-245 exhibited loss of steady state and reduced formate consumption.

To test whether complete carbon utilization could be achieved, the H_2_ fraction in the inlet gas stream was increased from 30 vol-% to 90 vol-% (period g). This led to an HTR_max_ of 17 mmol L^-1^ h^-1^, which equates to around four times more reducing equivalents supplied as H_2_ than as formate. Because, stoichiometrically, three moles of H_2_ are required for each mole of consumed formate, the condition should theoretically allow for full reduction of formate-derived CO_2_ to CH_4_. In other words, H_2_ supply could no longer be limiting because the minimum theoretical cH_2_ that could be maintained at full carbon utilization was 0.15 mM. However, the increased H_2_ supply negatively affected methanogenesis and formate metabolism of both strains (**Figure 2** – **period g**). The H_2_ uptake rate decreased and consequently cH_2_ increased.

Under these conditions, for *M. thermautotrophicus* Z-245, formate consumption ceased nearly instantaneous, and CO_2_ evolution dropped from 1.6 mmol L^1^ h^1^ to below 0.1 mmol L^-1^ h^-1^ (**Figure 2**). CH_4_ evolution showed a similar rapid decline. Although the culture partially recovered later in the cultivation period, the average formate uptake rate over the entire period was reduced by 80%. The *M. thermautotrophicus* ΔH (pFdh) culture initially responded more favorably to 90 vol-% H_2_. In the first hours after the switch, CH_4_ evolution increased to 80% carbon utilization. However, this improvement was transient, and we speculate that it might be related to dissolved inorganic carbon that was converted to CH_4_. The CH_4_ evolution could not be sustained and eventually decreased to rates lower than those observed before the switch in conditions. Formate uptake was also impaired, with an average reduction of 27% throughout the operating period (**Figure 2, Table 1** – **period g**). The bioreactor stabilized at a CH_4_ evolution rate of around 1.7 mmol L^-1^ h^-1^, despite the continued availability of CO_2_ and H_2_. Although *M. thermautotrophicus* ΔH (pFdh) tolerated the high H_2_ fraction better than *M. thermautotrophicus* Z-245, it also exhibited impaired methanogenesis and formate utilization under H_2_-excess conditions. At the end of the cultivation period, the formate concentration in the bioreactors were 31.2 mM and 81.7 for *M. thermautotrophicus* Z-245 and *M. thermautotrophicus* ΔH (pFdh), respectively. At these concentrations, the thermodynamic driving force favors the oxidation of formate (**Supplementary Figure S2, Equation S1, Text S1**).

In all tested conditions, *M. thermautotrophicus* Z-245 reached the highest stable carbon utilization of 36.3 ± 1.3% in period e (**Figure 1, Table 1**). *M. thermautotrophicus* ΔH (pFdh) sustained a stable formate utilization also in period f and reached a stable average carbon utilization of 63.4 ± 0.6% (**Figure 1, Table 1**).

### 3.2. Continuous cultivation with gas recirculation

Because increasing the H_2_ fraction from 30 vol-% to 90 vol-% did not improve carbon utilization and instead destabilized both cultures, we sought to enhance H_2_ availability without further increasing its fraction in the inlet gas. Therefore, after period g, we tried to recover the strains at a pH_2_ of 10 vol-% for two days before switching to 30 vol-% again. However, to increase gas-liquid mass transfer rate, we recirculated the off-gas through the bioreactor at the pH_2_ of 30 vol-%. This modification raised the estimated k_L_a by approximately one order of magnitude to 290 h^-1^ (not depicted). *M. thermautotrophicus* Z-245 was unable to recover from the prior exposure to 90 vol-% H_2_ and was, therefore, not extensively tested under gas recirculation conditions. The gas recirculation experiment was performed successfully only with *M. thermautotrophicus* ΔH (pFdh).

Formate uptake, H_2_ consumption, and CH_4_ evolution remained constant over the entire cultivation period with gas recirculation (Figure 3, Table 1 – period h). The average CH_4_ evolution rate during this period was 3.16 ± 0.12 mmol L^-1^ h^-1^, corresponding to a carbon utilization of 75.8 ± 4%, the highest stable value that we observed across all cultivation periods (Table 1 – period h). The estimated steady state cH_2_ was 0.12 mM, indicating that H_2_ was also not limiting under these conditions. Analogous calculations for dissolved inorganic carbon estimated a concentration of 1.8 mM. Despite the increased metabolic energy availability and higher CH_4_ production, the biomass concentration did not increase under gas recirculation conditions.

**Figure 3.**
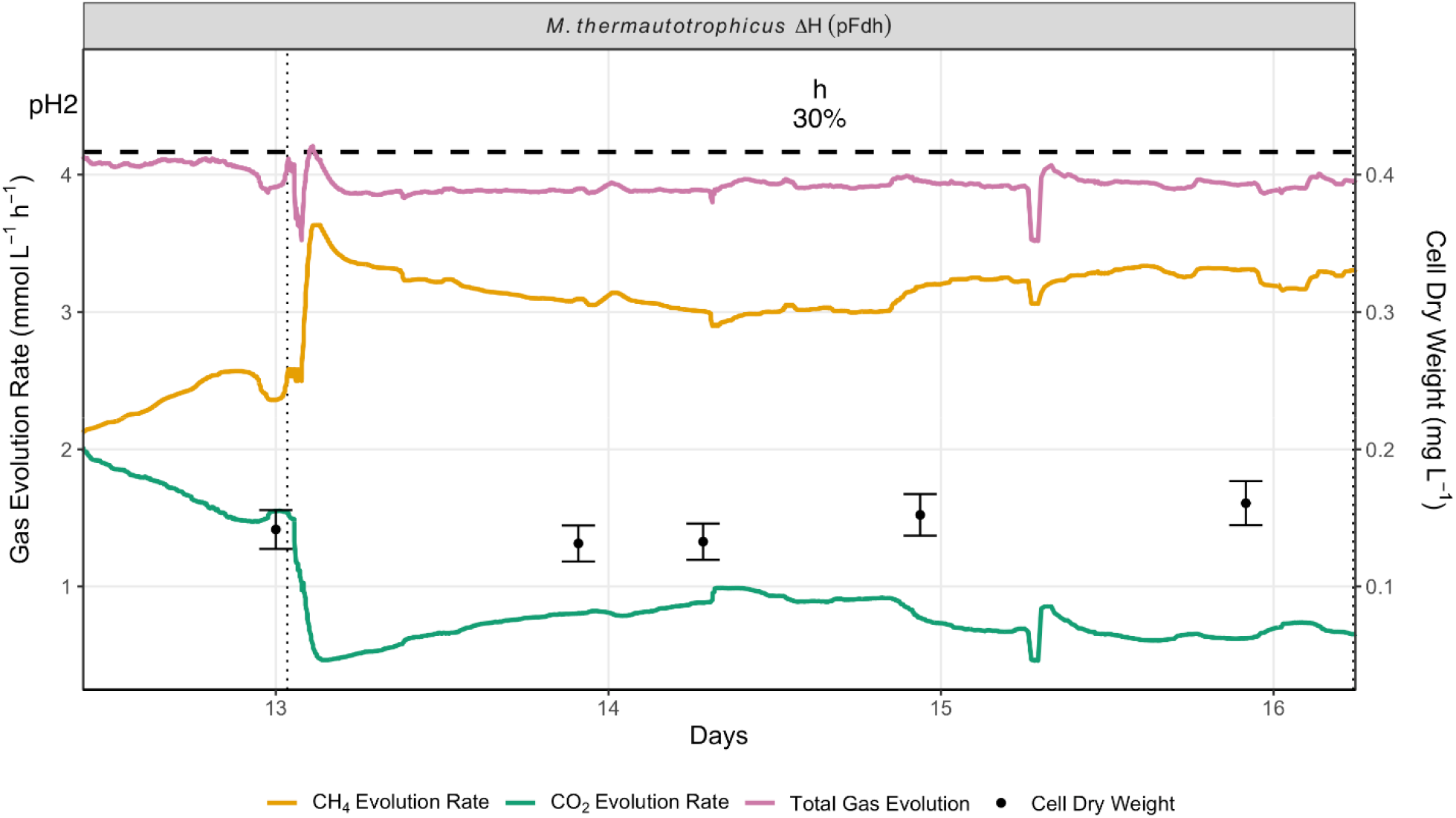
Gas evolution rates of M. thermautotrophicus ΔH (pFdh) under improved formate and H_2_ co-feeding conditions. After period g (90 vol-% H_2_) only M. thermautotrophicus ΔH (pFdh) was recovered successfully and further tested. The dotted black lines indicate the start and end of the gas recirculation period (period h). The H_2_ fraction of the inlet gas remained unchanged at 30 vol-%. Formate was fed at 4.16 mmol L^−1^ h^−1^ reflecting the carbon in flux and maximum theoretical carbon utilization (horizontal dashed line).

### 3.3. Biomass-specific production rates

With the stepwise increase in H_2_ supply, the CH_4_ evolution also increased. However, the biomass concentration in the bioreactor did not increase proportionally to the CH_4_ evolution. This is best observed in the CH_4_ biomass yield, Y_X/CH4_, defined as grams cell dry weight (gDW) biomass per mole of CH_4_ produced (**Table 1**). For *M. thermautotrophicus* ΔH (pFdh) grown on formate alone, Y_X/CH4_ was 1.6 gDW mol^-1^. Under gas recirculation at 30 vol-% H_2_, where cH_2_ was in excess, Y_X/CH4_ decreased to 0.95 gDW mol^-1^. This corresponds to an approximately 38% reduced biomass yield, indicating that under high H_2_ availability the culture became less efficient in converting the available reducing equivalents into biomass, pointing at other energy sinks or decreased catabolic yield. To test whether the system was nutrient limited, we conducted a serum bottle experiment on H_2_/CO_2_ (80:20 vol-%). Over the course of 6 days cells grew to 0.38 mg L^-1^ (**Supplementary Figure S4**). This was about twice as much as in the bioreactors, indicating that nutrients did not become limiting under the tested conditions in the bioreactors.

## 4. Discussion

Constitutive heterologous expression of an *fdh*-cassette from *M. thermautotrophicus* Z-245 in *M. thermautotrophicus* ΔH (pFdh) enabled continuous co-utilization of formate and H_2_ with a ∼ 2-fold higher stable conversion of the supplied carbon into CH_4_ compared to *M. thermautotrophicus* Z-245. The ability of *M. thermautotrophicus* ΔH (pFdh) to couple formate oxidation and CO_2_ reduction with H_2_, in a single continuous system, and to tolerate elevated H_2_ concentrations better than *M. thermautotrophicus* Z-245, highlights this strain as a promising biocatalyst for efficient formate valorization.

### 4.1. Physiological differences between *M. thermautotrophicus* Z-245 and *M. thermautotrophicus* ΔH (pFdh)

When formate and H_2_ contributed similar amounts of reducing equivalents (period – f), *M. thermautotrophicus* Z-245 showed a gradual decrease in formate uptake and CH_4_ production. We interpret this behavior as a regulatory or redox-controlled response, which could be linked to a switch in metabolism from F_420_-mediated methanogenesis to direct H_2_ utilization (Costa et al., 2013; Hendrickson et al., 2007; Reeve et al., 1997). Such a metabolic switch would be detrimental because CO_2_ from formate oxidation is the only carbon source and final electron acceptor under the applied conditions. *M. thermautotrophicus* ΔH (pFdh) showed no decline in formate uptake under the same conditions. This robustness likely reflects a reduced regulatory sensitivity, either due to intrinsic differences between the strains, or more importantly, due to the constitutive expression of the heterologous *fdh*-cassette, which bypasses native transcriptional control. However, the observed contrasting behaviors of the two strains at even higher pH_2_ cannot be fully explained by transcriptional regulation alone, as will be discussed independently in section 4.2.

Furthermore, during the initial cultivation on formate alone (*i*.*e*., without external H_2_), *M. thermautotrophicus* ΔH (pFdh) had a higher H_2_ fraction in the off-gas than *M. thermautotrophicus* Z-245 (1100 ppm *vs*. 200 ppm; **Supplementary Table S4 – period a**), consistent with results reported previously (Casini et al., 2023). The H_2_ release under formate-only growth suggests a difference in redox level between the two strains under identical growth conditions. Further studies are required to determine whether this behavior is a side-effect of constitutive expression of the *fdh*-cassette or reflects broader strain differences. Importantly, however, it indicates that *M. thermautotrophicus* ΔH (pFdh) is inclined to tolerate higher cH_2_ concentrations without a detrimental effect on formate consumption.

### 4.2. Catalytic limitations under excess H_2_ availability

When the H_2_ fraction was increased from 30 vol-% to 90 vol-%, cH_2_ shifted from near-zero to an estimated 0.4 mM, corresponding to a transition from H_2_ limitation to H_2_ excess. This switch proved problematic for *M. thermautotrophicus* Z-245 (**Figure 2**), as formate uptake and CH_4_ production collapsed directly. *M. thermautotrophicus* ΔH (pFdh) initially responded more favorably, but also showed a marked decline in formate and CH_4_ fluxes.

The timescale of the response rules out purely transcriptional regulation, which would be expected to be on a scale of hours instead of the observed response within seconds to minutes. Thus, it points towards catalytic or metabolic limitations directly imposed by the cellular redox state. For *M. thermautotrophicus*, the redox state of the F_420_ pool closely tracks H_2_ availability (de Poorter et al., 2005). At high H_2_ concentrations, the hydrogenases (*e*.*g*., FrhABG) rapidly reduce F_420_, whereas the re-oxidation rate depends strongly on the availability of the terminal electron acceptor CO_2_. Under excess H_2_ conditions, we expect the F_420_ pool to be largely in a reduced state (Xue et al., 2025). Fdh, however, needs oxidized F_420_ as an electron acceptor. A highly reduced F_420_ pool will, thus, directly inhibit formate oxidation, which likely explains the sharp decline in formate uptake and CO_2_ evolution observed immediately after the switch to 90 vol-% H_2_. The subsequent partial recovery of formate metabolism later in the cultivation period may be related to the increase in formate concentration in the bioreactor, which increases the thermodynamic driving force of the forward reaction (**Supplementary Figure S2**). The relatively higher affinity for F_420_ of Fdh (K_m_ ≈ 6 μM) compared to FrhABG (K_m_ ≈ 19 μM) (Jacobson et al., 1982; Schauer & Ferry, 1982), together with the possibility that Fdh can transfer electrons directly to heterodisulfide reductase, as suggested for other methanogens (Halim et al., 2024; Watanabe et al., 2021), may also prevent a complete shutdown of formate oxidation.

Interestingly, formate metabolism of *M. thermautotrophicus* ΔH (pFdh) was substantially less affected by the switch to 90 vol-% H_2_, compared to *M. thermautotrophicus* Z-245. This suggests that *M. thermautotrophicus* ΔH (pFdh) is intrinsically more robust to redox stress, potentially due to higher Fdh protein abundance arising from constitutive expression of the plasmid-encoded *fdh*-cassette. Future proteome analyses of Fdh-containing enzyme clusters formed under different redox conditions, may help elucidate whether electron bifurcation or competitive binding is employed. Overall, our results show that providing H_2_ in large excess does not lead to complete carbon utilization, but instead imposes catalytic and redox limitations that severely impair both formate oxidation and methanogenesis.

Matching the H_2_ supply to the stoichiometric and kinetic requirements of the metabolism will, therefore, be essential for an efficient bioprocess.

### 4.3. Maximizing carbon utilization with *M. thermautotrophicus* ΔH (pFdh)

Recirculation of the head-space increased the k_L_a by approximately one order of magnitude, which enhanced carbon utilization with *M. thermautotrophicus* ΔH (pFdh) without triggering the inhibitory effects that we observed at 90 vol-% H_2_. This demonstrates that high and stable co-utilization of formate and H_2_ is possible when H_2_ supply is optimized.

However, we did not achieve a carbon utilization above 75%. To explain this finding, we considered whether excess reducing power still imposed a redox-based inhibition of methanogenesis. However, because formate continued to be fully consumed under gas recirculation, we can rule out formate oxidation to be the primary limitation in this regime. Instead, improving gas-liquid mass transfer also increases stripping of dissolved gases, including CO_2_. At high mass-transfer rates, gas-liquid mass transfer removes CO_2_ that is produced from formate oxidation faster from the liquid, lowering dissolved inorganic carbon, and thereby limiting CO_2_ availability as the terminal electron acceptor for methanogenesis. Consistent with this, the off-gas CO_2_ fraction (∼ 1.5 vol-%) corresponds to a dissolved inorganic carbon level that can approach the reported K_m_ range (≈ 2 mM) for CO_2_/dissolved inorganic carbon-dependent methanogenesis in thermophilic systems (Xue et al., 2025). This suggests that CO_2_ stripping can constrain carbon utilization in bioreactors when gas-liquid mass transfer is optimized primarily for H_2_. Importantly, *M. thermautotrophicus* ΔH (pFdh) did not show the instability observed for *M. thermautotrophicus* Z-245 at elevated H_2_; instead, it maintained steady formate oxidation.

Bioreactor configurations designed specifically for gas fermentation, such as a bubble columns or trickle-bed bioreactors, could reduce CO_2_ stripping while maintaining high H_2_ transfer. Losses of carbon as CO_2_ are likely inherent due to the need for CH_4_ stripping. However, our results indicate that higher carbon utilization than the achieved 75.6% could be possible with *M. thermautotrophicus* ΔH (pFdh) without further genetic modification, if bioreactor design and operating conditions are tuned to balance H_2_ supply and CO_2_ retention.

### 4.4. Uncoupling of growth from methane production

Both strains showed a decrease in biomass yield with increasing H_2_ supply (**Table 1**). Serum bottle experiments confirmed that the observation was likely not related to nutrient limitations. The phenomenon of decoupling CH_4_ evolution and biomass production has been observed in methanogens before, and is often attributed to increased maintenance cost at higher methanation rates. However, why such uncoupling is condition-dependent and its underlying mechanisms remain poorly understood (de Poorter et al., 2007; Fardeau et al., 1987; Morgan et al., 1997). Furthermore, the observed decreased biomass yields at higher H_2_ availability in the presence of formate is different to observations that suggest an increased energy yield at cH_2_ above 0.08 mM for H_2_/CO_2_-only conditions (de Poorter et al., 2003).

Our data retain several constraints on mechanistic explanations. At 30 vol-% H_2_ with gas recirculation, *M. thermautotrophicus* ΔH(pFdh) reached a biomass-specific H_2_ uptake rate of 69.5 mmol_H2_ gDW^-1^ h^-1^, which is well below reported maximum H_2_ uptake rates of 200-400 mmol_H2_ gDW^-1^ h^-1^ for *M. thermautotrophicus* for H_2_/CO_2_-only conditions (de Poorter et al., 2005). Thus, the observed decrease in biomass yield from 1.6 to 0.95 gDW mol_CH4_^-1^ between formate-only and H_2_-excess conditions is unlikely to result from maximum biomass-specific uptake rates. From a physiological perspective, it is plausible that methanogens use CH_4_ production not only for ATP generation but also to maintain intracellular redox and ion gradients within a safe operating window. At high H_2_ availability, cells may need to increase CH_4_ flux to counteract the influx of reducing equivalents and prevent excessive reduction of intracellular cofactors. This might be especially important, when biomass synthesis cannot provide a sufficiently fast sink for ATP and reducing equivalents. Under such conditions, cells may respond by diverting more electron flow into CH_4_ formation by activating additional ATP-dissipating processes and/or the coupling efficiency between electron flow and ATP synthesis decreases under highly reduced conditions. The controlled growth conditions and real-time system characterization demonstrated in this work could be combined with high-resolution biomass composition analysis to help further elucidate inflexion points in biomass yields and biomass-specific rates.

### 4.5. Industrial implications

Industrial-scale bioreactors inevitably develop concentration gradients that can be substantial along the bioreactor height and across gas-liquid interfaces (Bokelmann et al., 2025). For formate-to-CH_4_ systems, the most consequential gradients are not only in formate, but in cH_2_, because local cH_2_ directly controls redox state and can shift methanogenesis from H_2_-limited to inhibition-prone regimes. This creates a scale-up challenge: even if the average cH_2_ is well controlled, local hot spots of high concentrations can occur near gas injection points, while H_2_ limitation can persist elsewhere.

Our results translate this challenge into a practical operating window. *M. thermautotrophicus* Z-245 began to deviate from stable, near-ideal behavior at relatively low dissolved H_2_ (∼ 0.016 mM), whereas *M. thermautotrophicus* ΔH (pFdh) sustained formate uptake and stable methanogenesis at ∼ 0.1 mM cH_2_. H_2_-excess conditions (cH_2_ ∼ 0.4 mM) impaired formate utilization and methanogenesis in both strains, but *M. thermautotrophicus* ΔH (pFdh) maintained higher activity and recovered quickly after lowering the cH_2_. Together, these observations suggest that *M. thermautotrophicus* ΔH (pFdh) offers a wider safe cH_2_ window, and therefore, greater process robustness under realistic mixing constraints, supporting higher overall rates, easier start-up, and more flexible operation during transients.

From a process-design perspective, the goal is typically an off-gas that is enriched in CH_4_ with minimal H_2_ and CO_2_. Achieving this, requires matching the H_2_ transfer rate to the formate-derived CO_2_ reduction demand, rather than simply increasing H_2_ partial pressure. A key constraint is the three-way coupling between gas-transfer objectives: CH_4_ must be stripped efficiently, H_2_ must be transferred into the liquid, and CO_2_ must be retained sufficiently to avoid electron-acceptor limitation. As a result, increasing k_L_a for H_2_ can unintentionally enhance CO_2_ stripping and cap achievable carbon utilization. Bioreactor configurations that are optimized for gas fermentation may offer a better balance by enabling high H_2_ transfer at lower superficial gas velocities and/or under elevated pressure while reducing CO_2_ losses. Operational levers, such as modest increases in pressure, gas recycle strategies, and pH/alkalinity adjustments to shift CO_2_ into the bicarbonate pool, could further improve dissolved inorganic carbon retention and move the system toward higher carbon utilization. Overall, *M. thermautotrophicus* ΔH (pFdh) appears particularly attractive for industrial application because its robustness at higher cH_2_ provides valuable headroom for dealing with unavoidable gradients while still maintaining stable formate conversion and methanogenesis.

### 4.6. Limitations and future directions

This study focuses on process-level physiology and does not directly resolve the molecular mechanisms underlying the observed differences between *M. thermautotrophicus* Z-245 and *M. thermautotrophicus* ΔH (pFdh). In particular, we infer regulation of formate dehydrogenase and redox constraints from gas fluxes and thermodynamic considerations rather than from direct measurements of Fdh abundance, activity, or its regulatory network. Likewise, while our data clearly show that elevated H_2_ partial pressures impair both formate metabolism and methanogenesis, we did not quantify how this relates to changes in the expression or post-translational state of key enzymes in the methanogenesis pathway. The plasmid in *M. thermautotrophicus* ΔH (pFdh) is essential for growth with formate. However, potential plasmid instability under H2-only transients in future applications could be overcome by integrating the *fdh*-genes into the chromosome of *M. thermautotrophicus* ΔH, while maintaining the constitutive expression. Future work combining the chemostat framework used here with targeted transcriptomics, proteomics, and enzyme activity assays will be important to clarify how Fdh is regulated in response to H_2_, and how *M. thermautotrophicus* ΔH (pFdh) and *M. thermautotrophicus* Z-245 differ in their metabolic adaptation to redox stresses.

## 5. Conclusion

This study demonstrates that constitutive expression of a heterologous formate dehydrogenase gene cassette enables stable co-utilization of formate and H_2_ in *M. thermautotrophicus* ΔH (pFdh), thereby significantly improving carbon utilization compared to *M. thermautotrophicus* Z-245 a native formatotroph. By decoupling formate metabolism from H_2_-dependent transcriptional regulation, *M. thermautotrophicus* ΔH (pFdh) sustained formate oxidation and CH_4_ production under conditions that inhibited *M. thermautotrophicus* Z-245, achieving up to 75.8% carbon conversion to CH_4_ in continuous cultivation. However, complete carbon utilization was not achieved, as excessive H_2_ availability introduced redox-related and catalytic limitations that impaired formate oxidation. These findings highlight that optimal process performance is not governed solely by substrate supply but also requires a balanced interplay between H_2_ transfer and intracellular redox state. Enhancing gas-liquid mass transfer *via* gas recirculation improved CH_4_ productivity without triggering inhibition, yet also revealed CO_2_ stripping as a potential bottleneck at high transfer rates. Overall, this work identifies *M. thermautotrophicus* ΔH (pFdh) as a robust and promising biocatalyst for formate-based biomethanation and provides important insights into the physiological and process constraints governing efficient carbon conversion. Future efforts should focus on optimizing bioreactor design and operating conditions to balance H_2_ supply and CO_2_ retention, as well as on elucidating the underlying metabolic mechanisms.

## 7. Data availability statement

All data supporting the findings of this study are available. A dataset containing utilized primers, the raw off-gas measurements, calculated gas evolution and uptake rates, and offline measurements, including the code used for processing online and offline data, and for generating all figures is available on Github (https://gitlab.com/stoutenlab/zipperle-hydrogen-formate-manuscript-2026).

## 8. Acknowledgments

This work was funded by the state of Baden-Württemberg (*Ministerium für Wissenschaft, Forschung und Kunst Baden-Württemberg*) and the CO_2_ Research Center funded by the Novo Nordisk Foundation with grant number NNF21SA0072700.4

