## Supplementary Material for "Towards complete carbon utilization: Improved methane yield from formate and hydrogen co-feeding through constitutive formate dehydrogenase-gene expression in *Methanothermobacter thermautotrophicus* ΔH"

*Table S1. Modified mineral salt medium composition (Balch 1979).*

| Component | per Litre | Final conc. |
|---|---|---|
| $NaHCO_2$ (sodium formate) | 13.600 g | 200.0 mM |
| $NaHCO_3$ (sodium bicarbonate)* | 6.000 g | 71.42 mM |
| NaCl | 0.450 g | 7.700 mM |
| $K_2HPO_4$ | 0.170 g | 976.0 µM |
| $KH_2PO_4$ | 0.230 g | 1.690 mM |
| $NH_4Cl$ | 0.500 g | 9.347 mM |
| $MgCl_2 \cdot 6H_2O$ | 0.080 g | 393.5 µM |
| $CaCl_2 \cdot 2H_2O$ | 0.060 g | 408.1 µM |
| $(NH_4)_2Ni(SO_4)_2$ — 0.2% w/v | 1 mL | 6.971 µM |
| $FeCl_2 \cdot 4H_2O$ — 0.2% w/v | 1 mL | 10.06 µM |
| Resazurin— 0.025% w/v | 4 mL | 4.363 µM |
| $Na_2MoO_4 \cdot 2H_2O$ | 0.00242 g | 10.00 µM |
| $Na_2SeO_3$ | 0.000173 g | 1.000 µM |
| L-Cysteine·HCl** | 0.500 g | 3.193 mM |
| Trace Element Solution | 1 mL | |
| Antifoarm 204*** | 0.06 mL | |

**Sodium bicarbonate was only added in media used in serum bottle cultivation as pH buffer.*

*** L-Cysteine hydrochloride was added anaerobically before autoclaving in serum bottles. For bioreactor runs, it was autoclaved separately from the medium and added before the start of the experiment to reduce the media.*

**** Antifoam was only added in media for bioreactors.*

*Table S2. Trace element solution (Casini 2023).The pH was not adjusted and the chemicals were added in the order in which they are listed.*

| Compound | | $g \cdot L^{-1}$ | Final conc. |
|---|---|---|---|
| Nitrilotriacetic acid (NTA) | $C_6H_9NO_6$ | 2 | 10.46 mM |
| Magnesium sulfate heptahydrate | $MgSO_4{\cdot}7H_2O$ | 30 | 121.72 mM |
| Manganese(II) sulfate | $MnSO_4$ | 5 | 33.13 mM |
| Sodium chloride | NaCl | 10 | 171.22 mM |
| Iron(II) sulfate heptahydrate | $FeSO_4{\cdot}7H_2O$ | 1 | 3.598 mM |
| Cobalt(II) chloride hexahydrate | $CoCl_2{\cdot}6H_2O$ | 1.8 | 7.563 mM |
| Calcium chloride dihydrate | $CaCl_2{\cdot}2H_2O$ | 1 | 6.807 mM |
| Zinc sulfate heptahydrate | $ZnSO_4{\cdot}7H_2O$ | 1.8 | 6.259 mM |
| Copper sulfate pentahydrate | $CuSO_4{\cdot}5H_2O$ | 0.1 | 400.5 µM |
| Aluminum potassium sulfate dodecahydrate (alum) | $KAl(SO_4)_2{\cdot}12H_2O$ | 0.18 | 379.4 µM |
| Boric acid | $H_3BO_3$ | 0.1 | 1.617 mM |
| Sodium molybdate dihydrate | $Na_2MoO_4{\cdot}2H_2O$ | 0.1 | 370.4 µM |
| Ammonium nickel(II) sulfate hexahydrate | $(NH_4)_2Ni(SO_4)_2{\cdot}6H_2O$ | 2.8 | 7.089 mM |
| Sodium tungstate dihydrate | $Na_2WO_4{\cdot}2H_2O$ | 0.1 | 303.2 µM |
| Sodium selenate | $Na_2SeO_4$ | 0.1 | 529.2 µM |

*Table S3. Primers used for strain identification (Casini 2023). The* fdh *operon in the genetically modified strain* M. thermautotrophicus *ΔH (pFdh) is encoded on plasmid pMVS1111A:*PhmtB-fdh$_{Z\text{-}245}$.

| Target | Forward Primer | Reverse Primer |
| --- | --- | --- |
| *Methanothermobacter thermautotrophicus* ΔH genomic DNA | ctgtccttatacctgctcctcctg | gatgcagatccctgcaacgg |
| *Methanothermobacter thermautotrophicus* Z-245 plasmid pFZ1 | ccaggaggtggaaaatgcatgc | gccgttttcatcaagcagcag |
| *Methanobacter marburgenisis* Marburg plasmid pME2001 | gttaatccagcacatcctcc | cctgtccaacttatacctttgg |

*Table S4. Average Fermentation data (± standard deviation) of each cultivation period for* M. thermautotrophicus *Z-245 and ΔH(pFdh). Gas recirculation was applied in condition h. $HTR_{max}$, maximal $H_2$ dissolving rate; HUR, $H_2$ uptake rate; CER, $CO_2$ evolution rate; MER, $CH_4$ evolution rate; FUR, formate uptake rate; OD, optical density; $cH_2$, dissolved $H_2$; CU, carbon utilization. For cultivation period g the last 2 hours were used for calculation.*

| Cultivation period | | a | | | | b | | | | c | | | | d | | | |
|---|---|---|---|---|---|---|---|---|---|---|---|---|---|---|---|---|---|
| pH2 | % | 0% | | | | 0.3% | | | | 1% | | | | 3% | | | |
| HTRmax* | mmol $L^{-1}$ $h^{-1}$ | 0.00 | | | | 0.06 | | | | 0.19 | | | | 0.57 | | | |
| HUR | mmol $L^{-1}$ $h^{-1}$ | 0.01 | 1.5% | -0.15 | 4.1% | 0.01 | 1.9% | -0.04 | 2.1% | 0.10 | 1.6% | 0.25 | 3.6% | 0.38 | 4.3% | 0.51 | 2.4% |
| CER | mmol $L^{-1}$ $h^{-1}$ | 2.97 | 1.4% | 3.10 | 3.3% | 3.00 | 1.8% | 3.06 | 2.0% | 2.97 | 1.5% | 2.99 | 3.6% | 2.92 | 4.1% | 2.98 | 2.2% |
| MER | mmol $L^{-1}$ $h^{-1}$ | 0.99 | 0.4% | 0.98 | 1.7% | 1.00 | 0.4% | 1.01 | 0.6% | 1.02 | 0.4% | 1.08 | 0.5% | 1.10 | 0.7% | 1.16 | 0.7% |
| FUR** | mmol $L^{-1}$ $h^{-1}$ | 3.97 | 1.5% | 4.08 | 3.8% | 4.00 | 1.9% | 4.07 | 2.1% | 4.00 | 1.5% | 4.06 | 3.6% | 4.03 | 4.2% | 4.14 | 2.3% |
| OD | - | 0.19 | 20% | 0.18 | 20% | 0.20 | 20% | 0.16 | 20% | 0.20 | 20% | 0.16 | 20% | 0.20 | 20% | 0.24 | 20% |
| cH2 | mM | 0.00 | 20% | 0.00 | 20% | 0.00 | 20% | 0.00 | 20% | 0.00 | 20% | 0.00 | 20% | 0.01 | 20% | 0.00 | 20% |
| Yx/CH4 | gDW $mol^{-1}$ | 1.66 | 20% | 1.60 | 14% | 1.73 | 20% | 1.44 | 20% | 1.73 | 20% | 1.33 | 20% | 1.63 | 20% | 1.83 | 20% |
| CU | % | 23.83 | 0.4% | 23.62 | 1.7% | 24.03 | 0.4% | 24.17 | 0.6% | 24.57 | 0.4% | 25.89 | 0.5% | 26.43 | 0.7% | 27.92 | 0.7% |
| e-balance*** | | 0.96 | 2.1% | 0.97 | 5.6% | 0.96 | 2.7% | 0.97 | 3.0% | 0.95 | 2.2% | 0.96 | 5.1%% | 0.94 | 6.0% | 0.99 | 3.3% |

| Cultivation period | | e | | | | f | | | | g | | | | h | |
|---|---|---|---|---|---|---|---|---|---|---|---|---|---|---|---|
| pH2 | % | 10% | | | | 30% | | | | 90% | | | | 30% | |
| HTRmax* | mmol $L^{-1}$ $h^{-1}$ | 1.91 | | | | 5.74 | | | | 17.20 | | | | 56.60 | |
| HUR | mmol $L^{-1}$ $h^{-1}$ | 2.08 | 4.8% | 1.76 | 2.9% | 5.26 | 6.3% | 6.53 | 3.5% | 3.05 | 12% | 3.73 | 3.8% | 8.73 | 18% |
| CER | mmol $L^{-1}$ $h^{-1}$ | 2.48 | 4.4% | 2.62 | 2.8% | 1.64 | 5.5% | 1.39 | 3.4% | 0.42 | 11% | 1.50 | 3.7% | 0.75 | 17% |
| MER | mmol $L^{-1}$ $h^{-1}$ | 1.52 | 1.3% | 1.46 | 0.5% | 2.30 | 2.2% | 2.64 | 0.6% | 1.16 | 3% | 1.74 | 0.4% | 3.16 | 4% |
| FUR** | mmol $L^{-1}$ $h^{-1}$ | 4.00 | 4.6% | 4.07 | 2.8% | 3.94 | 5.9% | 4.04 | 3.5% | 1.58 | 11% | 3.24 | 3.8% | 3.91 | 18% |
| OD | - | 0.22 | 20% | 0.21 | 20% | 0.30 | 20% | 0.38 | 20% | 0.23 | 20% | 0.34 | 20% | 0.34 | 20% |
| cH2 | mM | 0.00 | 21% | 0.01 | 20% | 0.02 | 21% | 0.00 | 20% | 0.48 | 23% | 0.46 | 20% | 0.10 | 27% |
| Yx/CH4 | gDW $mol^{-1}$ | 1.30 | 20% | 1.26 | 20% | 1.15 | 20% | 1.28 | 20% | NA | | NA | | 0.95 | 20% |
| CU | % | 36.45 | 1.3% | 34.97 | 0.5% | 65.90 | 2.2% | 63.38 | 0.6% | 73.3 | 3.4% | 57.1 | 0.5% | 75.83 | 4% |
| e-balance*** | | 1.01 | 6.6% | 1.02 | 4.0% | 0.98 | 8.6% | 0.96 | 5.0% | 0.97 | 16.5% | 0.96 | 5.3% | 0.95 | 25% |

** 20% standard deviation*

*** Disregards biomass as it is the sum of CER and MER*

**** Disregards biomass*

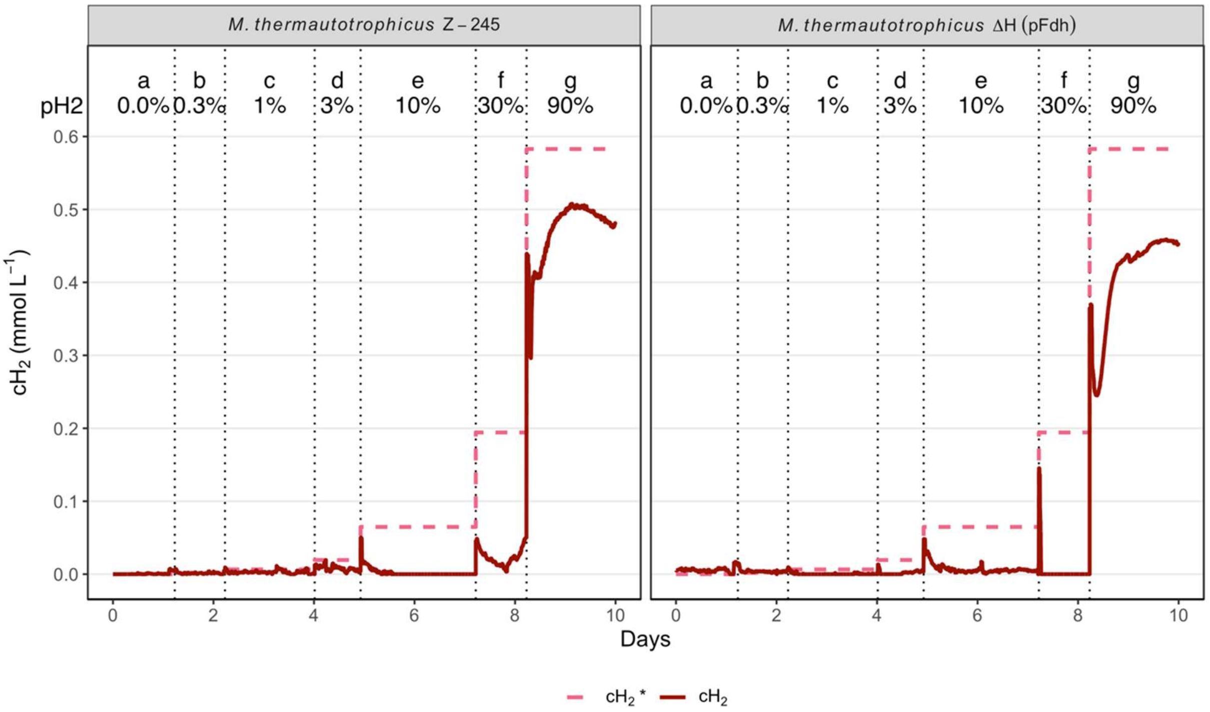


*Figure S1. Concentration of theoretical maximal dissolved $H_2$ concentrations (c*, rose) based on the partial pressure ($pH_2$) and calculated dissolved $H_2$ concentrations in the liquid phase ($cH_2$, red) over the different applied cultivation conditions. c* does not differ between strains, but $cH_2$ is influenced by the biological $H_2$ uptake rate, and thus, is different between the strains. Vertical dotted lines indicate a shift in cultivation conditions.*

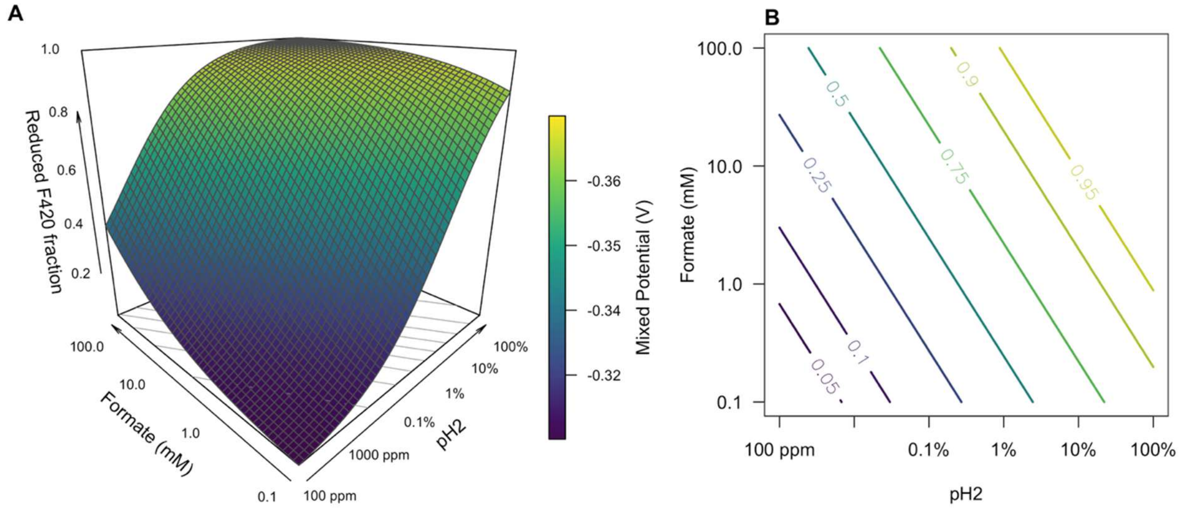


*Figure S2. Principle thermodynamic equilibrium analysis show that increased formate concentrations are required to allow oxidation through $F_{420}$ at increasing $cH_2$.* **(A)** *Log-log valley-plot for the $F_{420}$ redox fraction as a function of the $cH_2$ and formate concentrations.* ***(B)*** *A cut-through contour diagram at iso-redox conditions, for example, at a $pH_2$ of 90%, the equilibrium concentration $cH_2$ is 0.585 mM, and for 99% of the $F_{420}$ pool to be reduced (yellow iso-line), the formate concentration needs to be around 100 mM to maintain equilibrium.*

Equation $$E_{F_{420}} = E^0_{F_{420}} + \frac{RT}{2F}\ln\left(\frac{[F_{420}]}{[F_{420}H_2]}\right)$$

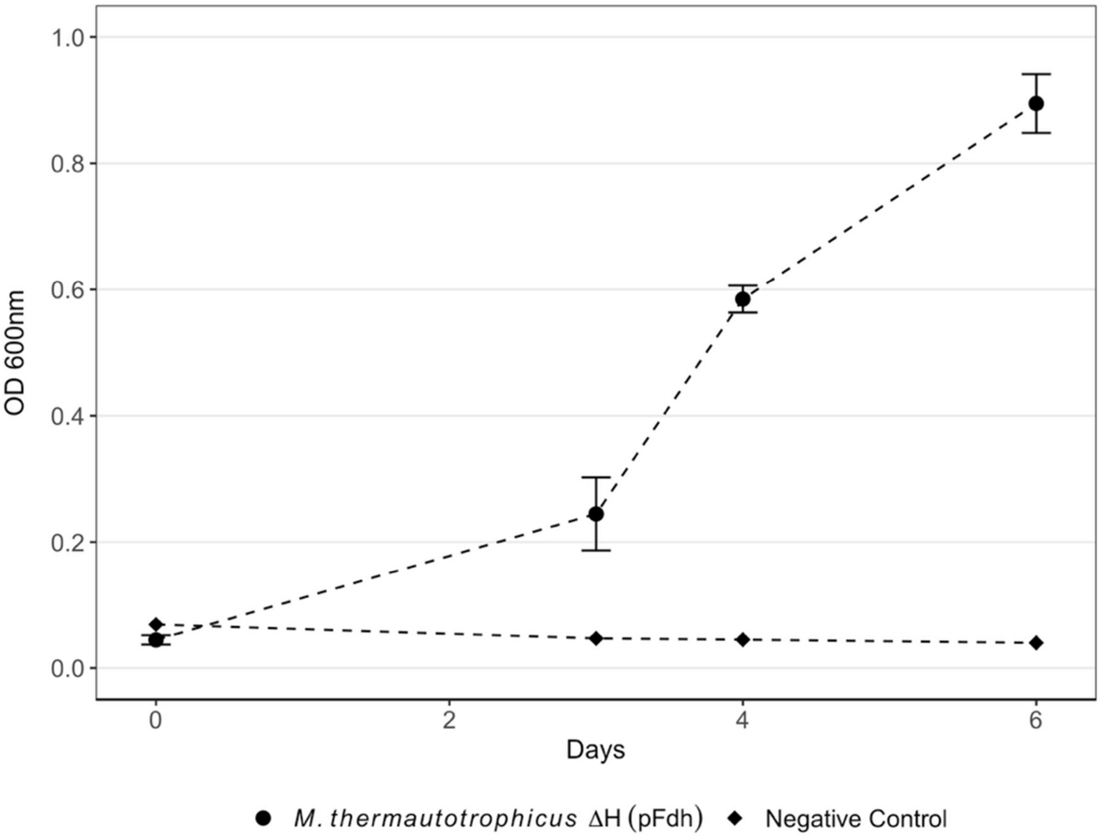


*Figure S3. Formate concentration in the bioreactors. Shown are condition f and g. The dotted line indicates the switch in condition. The bioreactor of* M. thermautotrophicus *Z-245 showed an increase in formate concentration in condition f while* M. thermautotrophicus *ΔH (pFdh) remained in chemostat. After the switch to condition g, both bioreactors accumulated formate indicating an impaired formate metabolism in both strains.*

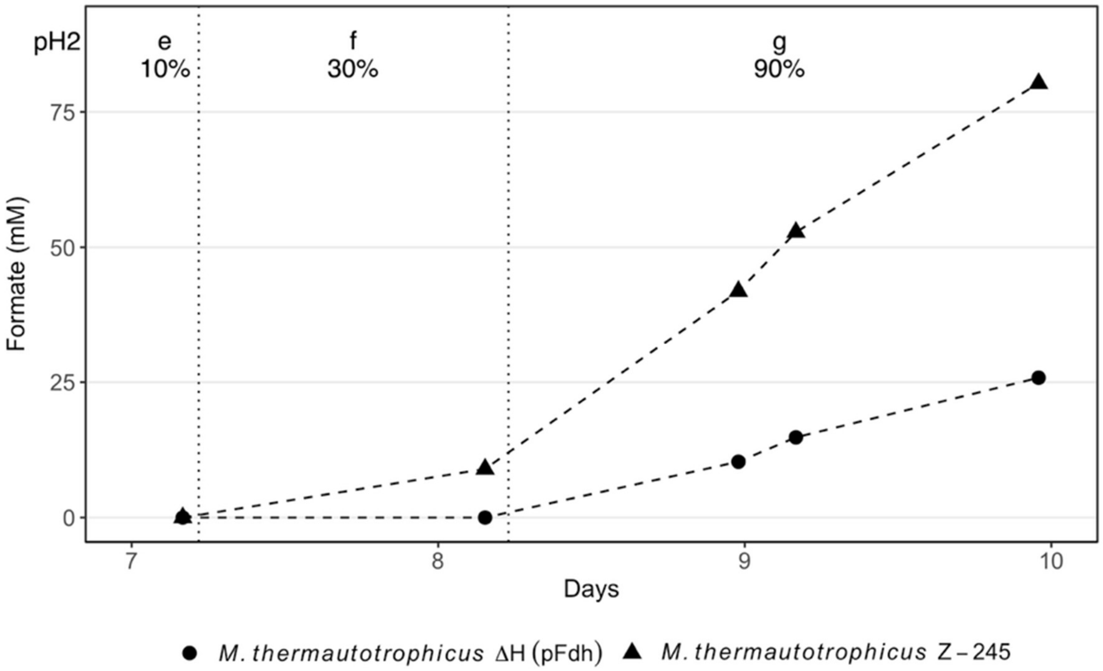


*Figure S4. Growth of* M. thermautotrophicus *ΔH (pFdh) on $H_2/CO_2$ (80:20 vol-%) in fed-batch. The error bars show the standard deviation. The serum bottles were replenished with fresh gas at least once per day (n=3).*

**Supplementary Text S1**

When multiple reversible redox couples supply electrons to the same carrier, the redox potential of the carrier is determined by a mixed potential defined by all participating couples (De Groot 2013, Dreyer 2016). In steady-state, both the $H_2/2H^+$ and $CO_2$/formate couples contribute thermodynamically to the $F_{420}/F_{420}H_2$ ratio. Enzyme affinity and catalytic efficiency do not alter the equilibrium redox potential but instead control the rate at which each donor drives the system toward that thermodynamically defined state and which donor dominates under non-equilibrium or substrate-limited conditions. Because $F_{420}$ is a finite cofactor pool, the $F_{420}/F_{420}H_2$ ratio is itself a function of the ambient redox potential. At high $H_2$ or formate levels, the pool becomes strongly reduced, depleting oxidized $F_{420}$. In this regime, the rates of both Frh- and Fdh-mediated electron transfer become limited by the availability of $F_{420}$, and their different $K_m$ values for $F_{420}$ begin to influence flux partitioning. Thus, while substrate affinities (for $H_2$ and formate) dominate at moderate redox poise, $F_{420}$-binding affinity becomes an important determinant of electron flux when the $F_{420}$ pool is driven toward full reduction. It is likely that cells employ alternative pathways to utilize formate when oxidation through $F_{420}$ is thermodynamically highly constrained. And equally likely is that downregulation of Fdh could help a cell retain electrons in the $F_{420}H_2$ pool. We can speculate that the absence of this regulation might allow *M. thermautotrophicus* ΔH (pFdh) to also mitigate redox stress by driving Fdh in the reverse direction and thereby produce formate.
